# Human gut Bacteroides uniformis utilizes mixed linked β- glucans via an alternative strategy

**DOI:** 10.1101/2021.06.24.449719

**Authors:** Ravindra Pal Singh, Raksha Thakur, Gulshan Kumar

## Abstract

Human gut microbial communities have a tremendous impact on well-being of the human health by producing myriad of metabolites. Polysaccharide utilizing capability of these microbial communities is a key driving force in shaping the composition of gut microbiota. Previous studies suggest that genes responsible for β- glucans utilizing in *Bacteroides* are present in a polysaccharide utilization locus (PUL). However, recent observations show that *Bacteroides uniformis* lacks such PUL archetypal organisation for efficient utilization of pustulan (β1-6) and mixed linkage (β1-4/ β1-3) glucans, and its mechanism is yet to be studied. Here, we first used *Bu*GH16 for production of important two mixed linked β- glucan-oligosaccharides, and then demonstrate kinetics of the *Bu*GH3_MLG_. The *B. uniformis* JCM13288^T^ coordinates between PUL associated glycoside hydrolase (GH)-16 and distantly localised bespoke GH 3 (*Bu*GH3_MLG_) for efficient utilization of mixed linkage (β1-4/ β1-3) glucans. *Bu*GH3_MLG_ found to be cleaved β1-4, β1-3 and β1-6 glucan-oligosaccharides. Such understanding of glycan utilization mechanism of human gut bacterial communities is crucial for development of nutraceutical therapy and improving health of human being.

## Introduction

Human gut microbiota (HGM) is an extraordinarily complex and dynamic community of microorganisms that are present in trillions, belonging to more than five thousand species (Almeida, *et al*., 2019). It is estimated that bacterial numbers of HGM is closer to human cells at a magnitude of 1:1 ratio (Sender, *et al*., 2016). Exploration of genomic sequences and metagenomic analysis of HGM is providing us a better clarification on taxon-level differences among microorganisms, and new metabolomic approaches are providing substantial understanding about metabolic fluxs between species (Sung, *et al*., 2017). Nevertheless, it restricts our amiability to fully underpin functional integrated information that can be applied to modulate HGM for improving the health of human beings. It is a well-known fact that the composition of HGM is linked to physiological condition of the gut. Metabolites of HGM, such as short chain fatty acids (SCFAs), are produced in the large intestine during microbial fermentation of indigestible glycans. SCFAs are the primary fuel source for colon epithelial cells (Topping, 1996), and protect from several gut-associated metabolic diseases including diabetes (Tan, *et al*., 2014). The main obstacle in manipulating the HGM dynamic for improving health of human beings is a paucity of understanding of the ecological forces that determine the form of community in individuals (Costello, *et al*., 2012, Heintz-Buschart & Wilmes, 2018). It is estimated that around 60 tonnes of food pass through the gastrointestinal tract in a human average lifetime (Azpiroz, 2005). These food contain complex dietary carbohydrates and the ability of the microbial communities to catabolise them is a key drivering factor in structuring HGM and its metabolic functions (Koropatkin, *et al*., 2012, David, *et al*., 2014). Therefore, underpinning glycan utilizing mechanism of each microbial species is needed to decipher their biological roles and development of nutraceutical intervention therapies to improve the health of human beings.

Gram-negative *Bacteroidetes* and Gram-positive *Firmicutes* of HGM are the predominant phyla in the colon of healthy individuals (Almeida, *et al*., 2019). *Bacteroidetes* are particularly known for utilizing a diverse range of dietary fiber by dedicating about 20% genes of their genomes (Martens, *et al*., 2011). These genes are present in a cluster of CAZymes, surface glycan-binding protein (SGBP), TonB-dependent transporter (SusC), and sensory/regulatory protein (hybrid two-component system), so-called polysaccharide utilization locus (PUL) (Lowe, *et al*., 2012). Generally, PUL encodes all components of proteins to complete saccharification of a specific glycan. For instance, a polysaccharide initially captured by the SGBP is broken down at the outer membrane of bacteria by an endo-acting enzyme(s) and then the released products (generally 2 to 8 degree of polymerization) are accessed by SusD. Afterwards, SucD makes a complex with SusC to import these products into periplasm for further digestion into their monomeric form by periplasmic enzyme(s), and then monosaccharides are released into the cytoplasm (Glenwright, *et al*., 2017). Such systems appear to be evolutionarily conserved among *Bacteroides* so as to prevent loss of monosaccharide in a highly competitive environment of the human gut. Complex molecular details of many PUL systems have been elucidated for pectin (Ndeh, *et al*., 2017, Luis, *et al*., 2018), yeast mannan (Cuskin, *et al*., 2015), xyloglucan (Larsbrink, *et al*., 2014), porphyran (Hehemann, *et al*., 2010), and β-glucans (Temple, *et al*., 2017).

*β*-glucans consist of linear and branched chain of *β*-D-glucose polysaccharides. It is naturally present in the cell wall of cereals, fungi, lichen, seaweeds, mushroom and bacteria with considerably varying physicochemical properties, depending on the source of origin (Rahar, *et al*., 2011). In particular, mixed linkage (*β*1-4/ *β*1-3) glucans are one of the main ingredients of routine diet and are commonly present in barley, lichenin and oat (Mikkelsen, *et al*., 2013). *β*-1-3 linked linear chain is present in bacterial cell wall (curdlan), and many *β*-1-3 linked linear chains are found to be having branches of a glucose (present in laminarin) or long chain of glucose (present in lentinan) by *β*-1-6 linkage(s). Many of these *β*-glucans have shown profound effects on the human immune system (Vetvicka, *et al*., 2019) and composition of the gut microbiota (Jayachandran, *et al*., 2018). Many of the bacterial taxa (including *Bacteroidetes*) present in HGM metabolize *β*-glucans and generate SCFAs for maintaining gut homeostasis (Singh, 2019).

Recently, our lab (Singh, *et al*., 2020) and others (Dejean, *et al*., 2020) have identified and dissected the molecular mechanism of the laminarin utilization locus (LUL) in the *Bacteroides uniformis* JCM13288^T^ and *Bacteroides uniformis* JCM5828 (ATCC 8492) respectively. These studies have established that LUL associated glycoside hydrolase-16 (GH16) can cleave mixed linkage (*β*1-4/ *β*1-3) glucan of the barley, but GH3 of the same PUL did not show any significant activity for digesting *β*1-4/ *β*1-3 linked oligosaccharides. This observation is far from the typical architecture of the characterized mixed linkage glycan utilization locus (MLGUL), reported from the *Bacteroides ovatus* ATCC8483 (Tamura, *et al*., 2017). The MLGUL contains bespoke enzymes for hydrolyzing barley polysaccharide at the outer surface and oligosaccharides in the periplasm. In addition, pustulan (β-1,6 linked glucan) utilization locus in the *B. uniformis* JCM13288^T^ also lacks exo-acting enzyme (GH3) as reported in the *B. thetaiotaomicron* VPI-5482.37, which is a part of fungal *β-*1-6 utilization locus (Temple, *et al*., 2017). To search the dedicated enzyme that can cleave *β*1-4/ *β*1-3/ *β-*1-6 in the *B. uniformis*, we used *B. uniformis* JCM13288^T^ as a model organism. We initially performed bioinformatic analysis for finding such candidate gene. Then, the putative gene was cloned and biochemically characterized.

## Material and methods

### General materials

*Bacteroides uniformis* JCM13288^T^ and *Bacteroides uniformis* JCM5828 (ATCC 8492) were purchased from Riken BRC Microbe Division (JCM). Strains were cultured under anaerobic conditions in Gifu anaerobic medium (GAM). All oligosaccharides were purchased from Megazyme (Ireland) or Sigma-Aldrich (India). AccuScript high-fidelity first-strand cDNA synthesis kit for reverse transcriptase-quantitative polymerase chain reaction (RT-qPCR) was procured from Agilent.

### Bacterial growth pattern and bioinformatics analysis

The growth curve pattern of the *B. uniformis* strains was monitored on 1% barley β glucan (BβG). Both strains were initially cultured in 5 ml GAM and once growth was seen, we performed growth curve experiment in minimal medium containing 1% BβG as per our previously used protocol (Singh, *et al*., 2020).

We recently identified two β glucans utilization loci in the *B. uniformis* JCM 13288^T^ and observed that PUL 36 (laminarin) associated *Bu*GH3_LM_ showed lower activity with gentiobiose (β-1, 6-linked disaccharide) and cellobiose (β1-4 linked disaccharide) (Singh, *et al*., 2020). It was also observed that *Bu*GH16 present within the LUL can digest mixed linkage (β1-4/ β1-3) glucan, so-called BβG (Dejean, *et al*., 2020). Overnight incubation of BβG with the *Bu*GH16 produced two limit digest products Glu-β-1-4-Glc-β-1-4-Glc-β-1-3-Glc (G4G4G3G) and Glc-β-1-4-Glc-β-1-3-Glc (G4G3G or cellobiose-glucose). Given the lower activity of *Bu*GH3_LM_ toward cellobiose, it was predicted that there might be another bespoke enzyme produced by the *B. uniformis* for efficient cleavage of limit digest products in the periplasmic space. Therefore, we revisited the genome of the *B. uniformis* JCM 13288^T^ searching for an exo-acting enzyme that may show comparatively high activity in periplasmic space against gentiobiose, G4G4G3G, G4G3G and cellobiose. All 33 GH3 were annotated using the CAZymes pipeline and out of them, 3 genes (BUNIF_00608, BUNIF_04508 and BUNIF_04929) showed 74, 58, and 58 % homologous to BT3314, respectively. The BT3314 (GH3) of the *B. thetaiotaomicron* VPI-5482.37 was identified as a part of the fungal β-1,6 glucan utilization locus (Temple, *et al*., 2017). LipoP 1.0, PSORTb and SignalP 5.0 servers were used to predict the lapidated signal peptide sequence and cellular localization of the BUNIF_00608, BUNIF_04508 and BUNIF_04929.

### Total RNA extraction and Gene expression analysis

Changes in the levels of transcript expression of selected three genes of the *B. uniformis* JCM 13288^T^ were done by RT-qPCR. The strain was initially cultured in 5 ml of GAM broth for 48 h, and then cells were spin down by centrifugation (6000 × g), followed by washing thrice with autoclaved phosphate-buffered saline (PBS). 20 µl of 1 ml suspension of cells was inoculated in 10 ml minimal medium containing 1% (w/v) laminarin, BβG or lentinan. About 3-5 ml of bacterial suspension were harvested at the mid-log phase (A_600_ about 0.6) by centrifugation at 15000 x g, and cell pellets were resuspended in 300 µl of a solution consisting of 0.02 M sodium acetate (pH 5.5), 0.5% SDS, and 1 mM ethylenediaminetetraacetic acid (EDTA). Right after that, 300 µl of phenol (saturated with citrate buffer, pH 4.3, P4682-Sigma Aldrich) was added, and the solution was incubated at 60°C for 5 min with mild shaking. It was then centrifuged at 15000 x g for 5 min and the upper aqueous phase was transferred into a new micro-centrifuge tube. The obtained upper aqueous phase was mixed with three volumes of chilled ethanol and then left for 30 min at -80 °C. Followed by, the precipitate was collected by centrifugation at same speed, and pellet containing RNA was serially washed twice with 70% and 100% ethanol. Obtained final white pellet was dissolved in 200 µl of distilled water for further processing. The concentration of obtained total RNA was determined by NanoDrop™ (Thermo Scientific). DNase 1 treatment was also given if residual DNA was observed, following the manufacturer’s instructions (Sigma Aldrich-AMPD1-1KT).

About 2 µg of obtained total RNA was immediately used for cDNA formation using AccuScript High Fidelity cDNA Synthesis Kit (Agilent). RT-qPCR was performed in a 96-well plate on a CFX96 qPCR system (BioRad), with SYBR® Green Master (BioRad) mix and specific primers as mentioned in supplemental Table S1. The qPCR reactions were performed in 10 µl batch, consisting of 5 µl of SYBR Green mix, 30 ng of cDNA, and 1 pmol of each primer of a gene or 16S rRNA. The reaction condition was 95 °C for 10 min, followed by 40 cycles at 95 °C for 10 s, 55 °C for 10 s, and 72 °C for 10 s. The ΔΔC_T_ method was used for calculating changes in transcript expression of each gene, and data were normalized to 16 S rRNA transcript level. Finally, changes in the transcript expression level of *bUNIF*_*00608, bUNIF_04508* and *bUNIF*_*04929* were represented as fold change, in comparison between transcript expression levels of these genes in minimal medium containing glucose versus specific carbon source.

### Cloning, expression, and purification of enzymes

Based on the gene expression profile, the nucleotide sequence of a highly transcribed gene, BUNIF_04929, was retrieved from the genome of *B. uniformis* JCM 13288^T^, and appropriate primers were designed (supplementary Table 1) after excluding lapidated and signal peptide sequence (Juncker, *et al*., 2003). Gene was amplified using a high-fidelity polymerase with primers specifically designed to perform Gibson assembly in *BamH* I and *Xho* I digested pET28a vector (Novagen), to express gene with N-terminal (His) _6_-tag. A cloned construct was initially transformed into the *Escherichia coli* TOP10 cells. A positive clone was selected from Petri-plates and then plasmid from positive clone was extracted using the GeneJET Plasmid Miniprep Kit (Thermo Fisher Scientific). The integrity of cloned gene in the pET28a in terms of orientation and sequence was checked at this stage, and it was then transformed into *E. coli* BL21 (DE3) (Rosetta, Novagen).

For expression of proteins, Rosetta cells containing desired gene were cultured in Luria–Bertani (LB) medium with chloramphenicol (35 µg / ml) and kanamycin (50 µg / ml) to OD ∼ 0.6 at 37 °C. Afterwards, expression of the protein was induced by adding 0.5 mM isopropyl β-D-1-thiogalactopyranoside. It was further incubated at 18 °C for 16 h. Post incubation, bacterial cells were harvested by centrifugation at 4,000 x g and then re-suspended in phosphate buffer (50 mM, 300 mM NaCl, pH 7.4) with an appropriate EDTA-free protease inhibitor cocktail (Sigma Aldrich, Cat. No. 4693159001). Cells were lysed by ultrasonication, and supernatant was collected by centrifugation at 16,000 x g for 30 min at 4 °C. Supernatant was initially concentrated through Amicon® Ultra-15 Centrifugal Filter Unit (10 kDa) and then about 5 ml of it was mixed with 20 ml of the same phosphate buffer, containing a suspension of nickel-nitrilotriacetate resin. It was then incubated for 30 min at 4 °C. Followed by, the resin was filled in column and one-step purification was carried out using elution buffer (50 mM phosphate buffer, 300 mM NaCl, 150 mM imidazole, pH 7.4) to get purified protein. Fractions containing targeted proteins were pooled, and then residual imidazole was exchanged with 50 mM HEPES buffer by Amicon® Ultra-15 Centrifugal Filter Unit (10 kDa). A 10 % sodium dodecyl sulfate-polyacrylamide gel electrophoresis (SDS-PAGE) was performed for determining molecular weight of the desired protein. The concentration of purified protein (we named as *Bu*GH3_MLG_) was finally determined by NanoDrop™, and then protein was stored at -80°C, having a concentration of 1.2 mg/ml for biochemical analysis.

### Evaluating biochemical properties of enzymes

Optimum pH for *Bu*GH3_MLG_ was determined by incubating the reaction mixture at 37°C for 10 min in a variety of buffers (50 mM), and p-nitrophenyl-β-laminaribioside (pNP-Laminaribiose) is used as a substrate. A standard reaction mixture of 100 µl volumes, consisting of 10 µl of 10 mM pNP-laminaribiose, 10 µl of 1.2 mg/ml of *Bu*GH3_MLG_ and 80µl of different buffers (50 mM) were used. Different buffers such as citrate (pH 3 to 5.5), sodium – phosphate (pH 5.7 to 8), sodium acetate (pH 4.5 to 6), Tris-HCl (pH 8 to 10) and HEPES (pH to 5.6 to 7.4) were used. For determining the optimal temperature of the *Bu*GH3_MLG_, standard reaction volume of 100 µl having 10 µl enzyme (1.2 mg/ml) and 10 µl of 10 mM of pNP-laminaribiose was performed at various temperatures (15, 20, 25, 30, 37, 48 and 55 °C) for 10 min using optimized pH. All reactions were ended by adding 2 volumes of 1 M Na_2_CO_3_ and the amount of released p-nitrophenol was calculated by measuring the absorbance at 410 nm using NUNC 96 well plates on a SpectraMax® i3x Multi-Mode Microplate Reader. Impact of metals and detergents on *Bu*GH3_MLG_ activity was determined by taking 2 mM concentration of different salts such as MnCl_2_, MgCl_2_, CaCl_2_, FeSO_4_, MgSO_4_, LiBr, NaNO_3_, NH_4_Cl, KCl, CoCl_2_, NiSO_4_, NaCl, and detergents like urea, 1,4-dithiothreitol (DTT), and EDTA. All these reactions were performed in a standard volume mixture in sodium phosphate buffer (pH 5.7) buffer, and buffer with no added supplement was used as control.

The capability of *Bu*GH3_MLG_ for degrading various oligosaccharides (like, cellobiose, cellotriose, cellotetraose, laminaribiose, laminaritriose, laminaritetrose, gentiobiose and other disaccharides) was screened out in 100 µl reaction solution, containing 10 µl 100 mM substrate and 10 µl enzyme (1.2 mg/ml) in 100 mM sodium phosphate buffer (pH 5.7) for 4 h at 37 °C, and overnight to understand limit degraded products. After incubation, the reaction was stopped by heating at 100 °C (5 min) and degraded products of oligosaccharides were analyzed by thin-layer chromatography (TLC). TLC analysis was performed on Silica Gel 60 F254 (Merck) using butanol: ethanol: water (5:3:2, v/v/v) mobile phase, and produced sugars were visualized by spraying TLC with 5% H_2_SO_4_ in ethanol, followed by heating.

The Michaelis–Menten kinetics in terms of K_m_, K_cat_ and K_cat_ / K_m_ of the *Bu*GH3_MLG_ were determined using a range of concentrations of various p-Nitrophenyl (pNP)-β-D-sugars. Reactions were performed in 100 µl standard reaction volume as used for determining hydrolysing capability of oligosaccharides. Termination and reading of all reactions were carried out as mentioned above.

For estimating kinetic parameters of BPGH94-2 using PAHBAH reducing sugar assay, calibration was conducted by hydrolysis different concentrations (0, 2, 4, 8, 16, 32, 64, 100, 120, 140, 160 and 180 µM) of oligosaccharides for 30 min. 150 µl of a freshly prepared 8:2 mixture of reagent A (50 mM trisodium citrate, 10 mM CaCl2, 0.5 M NaOH) and reagent B (50 mM sodium sulphite, 0.3 M 4-hydroxybenzhydrazide and 0.6 M HCl) was added to 50 µl of sample. The reaction cocktail was boiled for 10 min at 100°C. Once reaction was cooled down and absorbance was taken at 410 nm using the SpectraMax® i3x Multi-Mode Microplate Reader. In order to calculate the amount of released glucose reducing end equivalents, a calibration curve with glucose (0, 2, 4, 8, 16, 40, 80, 100, 200, 300 and 500 µM) was used for every experiment. Same concentrations of samples were also used for control measurement and value of control was subtracted from test reading before calculating kinetics parameters.

### Production of limit degraded products of barley β glucan (BβG) by BuGH16, purification and analysis

As both strains (*B. uniformis* JCM 5828 and JCM 13288^T^) showed growth on mixed linkage BβG, we explored the possible role of *Bu*GH3_MLG_ in utilizing oligosaccharides generated by *Bu*GH16. The *Bu*GH16 of the *B. uniformis* JCM 13288^T^ showed 100% homology to one protein characterized from *B. uniformis* JCM 5828 (Singh, *et al*., 2020). A study by Dejean, *et al*. (2020) observed that *Bu*GH16 of the *B. uniformis* JCM 5828 can digest BβG and produced two mixed linkage compounds, G4G4G3G and G4G3G.

Therefore, we used *Bu*GH16 of the *B. uniformis* JCM 13288^T^ for generating limit-digest products from BβG by incubating the reactions overnight at 37°C using multiple replicates, each of 100 µl volume, having 10 µl of enzyme (1 mg/ml) and substrate (10 mg/ml) in sodium – phosphate buffer (pH 5.7) in each batch. The next day, reaction mixture was terminated by heating and limit-digested products were recovered by doing centrifugation (10000 × g) for 10 min and then applied to Toyopearl S40 resin containing column (GE Healthcare-XK 26/100) for purification. All fractions eluted from the column were initially viewed using TLC and similar fractions were pooled together; leading to production of two major fractions. Both fractions were then freeze-dried. MALDI-TOF and ^13^C-DEPT NMR were performed as carried out previously (Singh, *et al*., 2020). MestreNova software was used for ^13^C-DEPT NMR data processing.

### Sequence alignment and phylogeny analysis

Amino acid sequence of *Bu*GH3_MLG_ was used to develop a homology model using the Phyre2 platform. Afterwards, amino acid sequences of structurally characterized GH3 were retrieved from the protein data bank (RCSB-PDB) and used for deciphering their evolutionary relationship via molecular evolutionary genetics analysis X (MEGA X) software. The evolutionary phylogenetic tree of those sequences was reconstructed using the Neighbor-Joining method through the bootstrap test. Structure-based sequence alignment of the BuGH3_MLG_, BoGH3_MLG_ and BT3314 (GH3) with structurally characterized GH3 β-glycosidase was performed using the ESPript 3.0 (Robert & Gouet, 2014).

## Results and discussion

### Bacterial growth on mixed linkage β glucans and gene expression

Both strains showed robust growth on BβG (Fig.1A). Cereal β glucans (such as barley) are one of the most abundant components of soluble dietary fibers, and have great nutritional and functional properties (Turck, *et al*., 2021) with positive modulatory effect on gut microbiota (Hughes, *et al*., 2008). BβG consists of a short stretch of β-1,4 linked chain connected with β-1,3 linked glucose units (Izydorczyk & Dexter, 2008, Mikkelsen, *et al*., 2013). To digest such mixed linkage glucans, human gut bacteria encode a cluster of genes in the form of PUL. Such a PUL is well characterized in *Bacteroides ovatus* ATCC 8483 (Tamura, *et al*., 2017). Both *B. uniformis* strains do not have PUL for mixed glucan utilization; however, they have PUL for utilizing laminarin (Singh, *et al*., 2020). On the contrary, *B. ovatus* ATCC 8483 does not possess homologous LUL, and thus, does not grow on laminarin (Martens, *et al*., 2011). Given these interesting facts, three GH3 of the *B. uniformis* 13288^T^, based on high percentage homology to BT3314, were selected with the prediction that they may degrade β-1-4 and β-1-6 of generated oligosaccharides from BβG and pustulan. Gene expression profiling of those indeed favoured this prediction where *bunif_04929* was overexpressed about 805, 455, and 5 fold when the bacterium was grown in minimal medium containing laminarin, BβG and lentinan as compared to glucose respectively. The *bunif_00608* was also overexpressed as compared to glucose with 10, 6 and 5 fold when the bacterium was grown in minimal medium containing laminarin, BβG and lentinan respectively (Fig.1C). It was demonstrated that PUL associated components such as SusC and SGBPs were significantly upregulated when their transcriptional expression was measured in the presence of BβG as the sole carbon source (Dejean, *et al*., 2020). This observation highlights that *bunif_04929* may have role in BβG utilizing, therefore, it was cloned and its functional characteristics were performed. LipoP 1.0 (Juncker, *et al*., 2003), PSORTb3.0 (Yu, *et al*., 2010) and SignalP 5.0 (Almagro Armenteros, *et al*., 2019) servers indicated that BUNIF_00608, BUNIF_04508 and BUNIF_04929 have a lapidated signal peptide sequence type 1 for periplasmic space (PSORTb score 10.00).

**Fig. 1.**
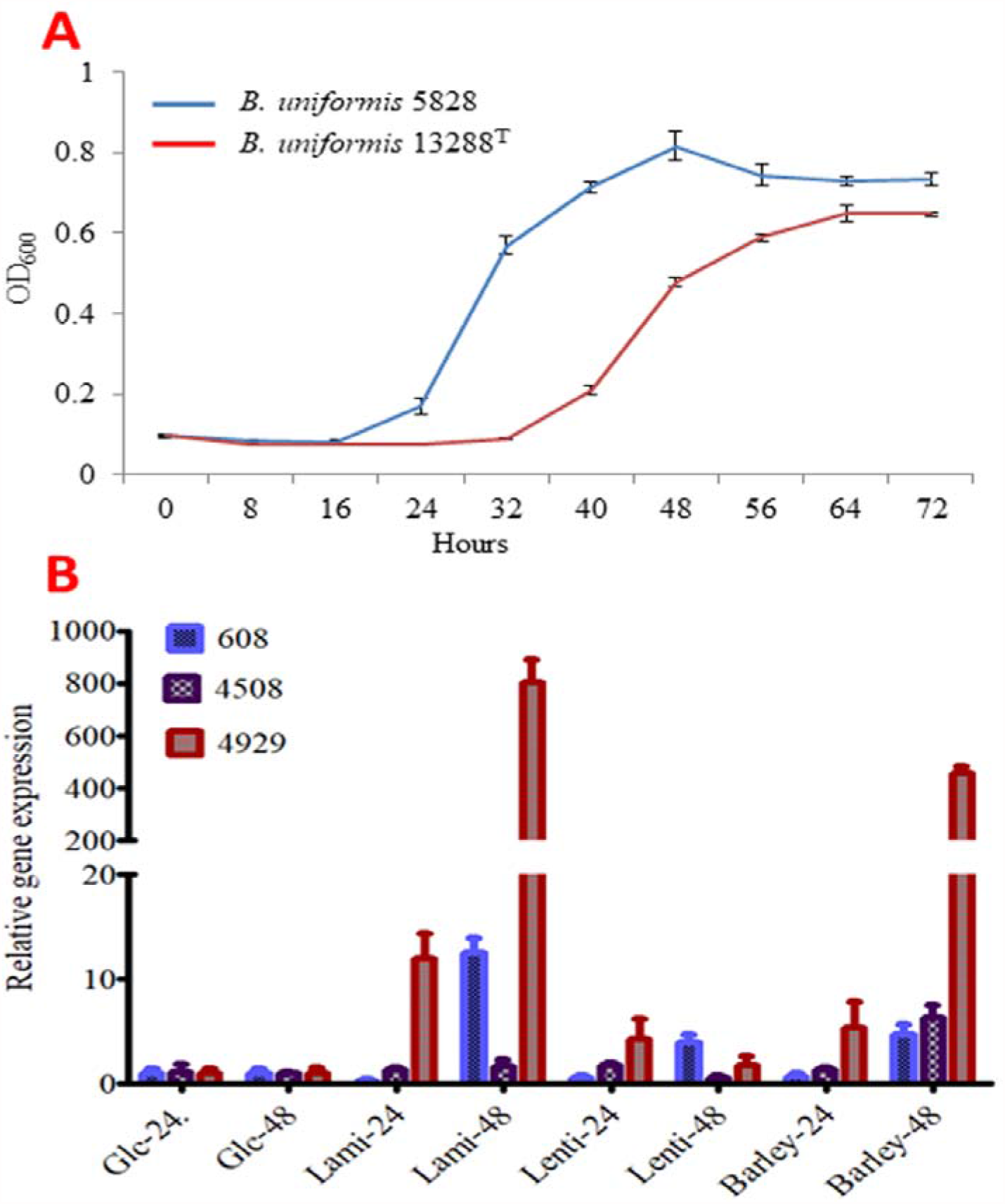
Bacterial growth patterns, and reverse transcription-quantitative polymerase chain reaction (RT-qPCR). (**A**) Growth pattern of both *Bacteroides uniformis* strains on 1% barley β glucan. (**B**) Gene expression of selected genes when bacterium was grown on minimal medium having 1% laminarin, lentinan or barley β glucan. Fold change in transcript expression of selected genes reported as defined polysaccharide versus glucose.

### Heterologous expression of BuGH3_MLG_, and pH, temperature, metal ion optimization

The *bunif_04929* was cloned into pET28a with a strategy to have a hexahistidine tag at the N-terminus of the protein. Subsequently, nickel affinity chromatography yielded about 25 mg per litre of purified Bunif_04929 and was concentrated to 1.2 mg/ml, subsequently stored at -80 °C until required. About 10 µg of the protein was run on the 10 % SDS-PAGE and the band was observed to be around 75 kDa (Fig. 2A) as it was predicted from the total number of amino acids of the protein including hexahistidine tag.

**Fig. 2.**
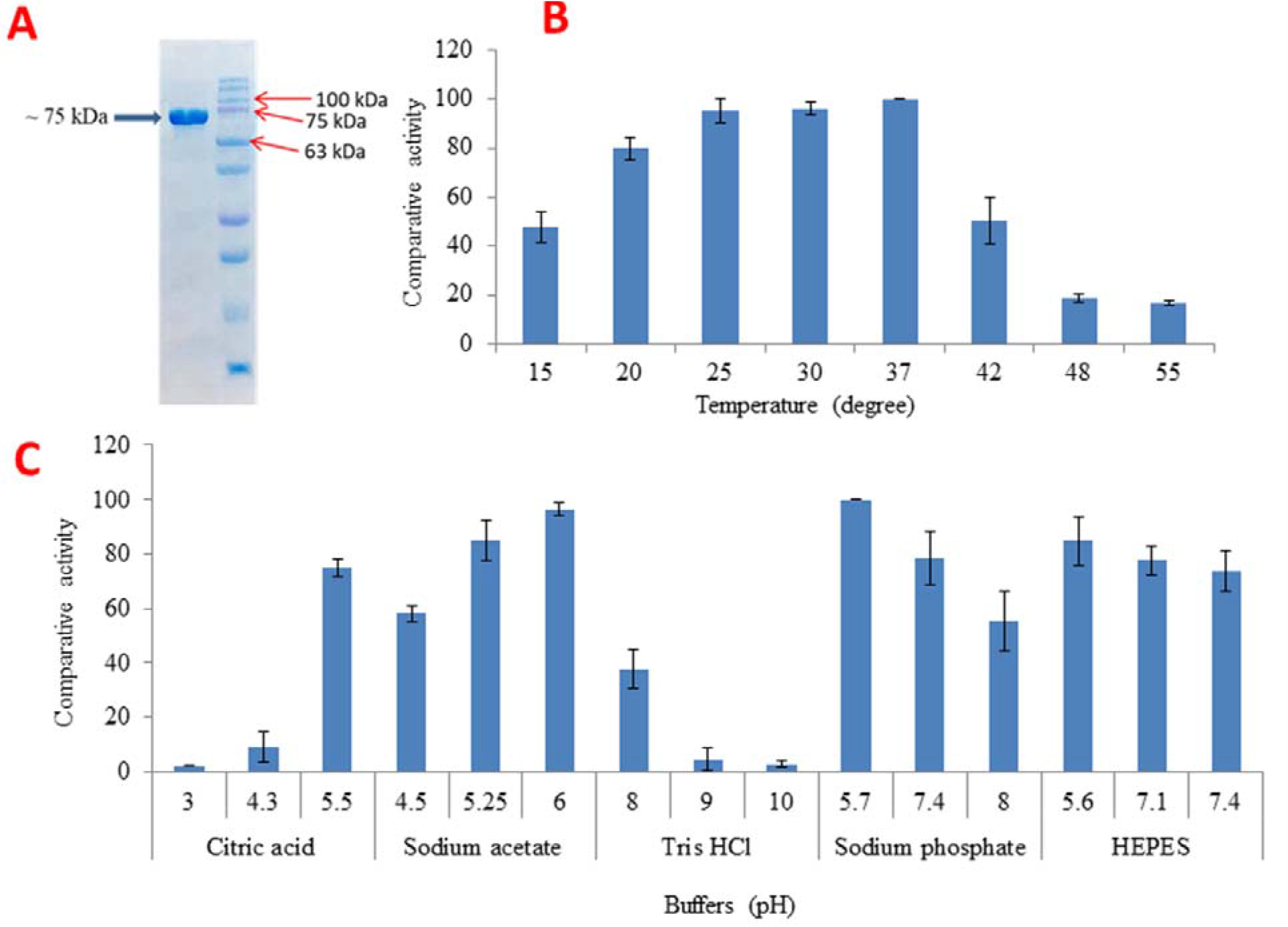
Sodium dodecyl sulphate-polyacrylamide gel electrophoresis (SDS-PAGE) analysis and optimization condition of the *Bu*GH3_MLG_. (**A**) The 10% SDS-PAGE was used to estimate molecular weight of recombinant protein. Different ranges of temperatures (**B**) and buffer systems (**C**) were used to optimize conditions of enzyme activity.

It showed more than 80% residual activity at a range from 20 to 30 °C with highest at 37 °C (Fig. 2B). The activity was drastically reduced below 20 and above 42 °C. Interestingly, it showed more than 50% activity from pH 4.5 to 7.4 (Fig. 2B), at different buffers. Given the broad activity range of *Bu*GH3_MLG_ on different buffers, we decided to check the impact of metal ion on its activity. A wide variety of selected metals (2 mM concentration) did not significantly influence the activity of *Bu*GH3_MLG_ (Table S2) whereas urea, DTT and EDTA drastically minimized the activity, reflecting that metal might be playing an important role in catalytic activity at higher concentration. Indeed, increasing the concentration of NaCl to 5 and 10 mM profoundly boosted the enzymatic activity.

### Identification of BβG products by BuGH16

The *Bu*GH16 yielded majorly two compounds after overnight incubation with BβG (Fig. 3A) that were eventually separated by Toyopearl S40 resin. Molecular masses of both compounds were found to be 689.19 [M + Na]^+^ and 527.17 [M + Na]^+^ when they were analyzed with high-resolution MS using MALDI-TOF (Fig. 3 B and C). Purified compounds were freeze-dried and glycosidic bonds were characterized with ^13^C-DEPT NMR. The ^13^C-DEPT NMR allowed us to determine anomeric carbon of non-reducing, middle and reducing end residues and the presence of glycosidic bonds. Some characteristic signals at ca. δ 78.0 belong to C-4 originated from non-reducing end glucose (Singh, *et al*., 2020). The presence of one and two signals at ca. δ 78.0 indicate that 527.17 [M + Na]^+^ and 689.19 [M + Na]^+^ have one and two C-4 linked glucose units respectively (Fig. 3 D and E). Presence of ca δ 84 and – 82 signals indicate the presence of β-C-3 and α-C-3 of glucopyranose bearing glycosyl residue at reducing end (Singh, *et al*., 2020). Therefore, 689.19 [M + Na]^+^ compound was identified as G4G4G3G [β-d-Glc^iv^(1→4)β-d-Glc^iii^(1→4)β-d-Glc^ii^(1→3)-d-Glc^i^] with partial assignment through ^13^C NMR (101 MHz, D_2_O) δ 102.84 (C-1^iv^ β), 102.60 (C-1^iii^ β), 102.28 (C-1^ii^ β), 102.27 (C-1^ii^ α), 95.61 (C-1^i^ β), 91.97 (C-1^i^ α), 84.34 (C-3^i^ β), 82.10 (C-3^i^ α), 78.29 (C-4^ii^ β), 77.94 (C-4^iii^ β), 75.47, 75.27, 74.74, 74.01, 73.74, 73.14, 72.75, 72.40, 71.10, 70.85, 68.47, 68.00, 61.00, 60.94, 60.60, 59.83. The 527.17 [M + Na]^+^ compound was identified as G4G3G [β-d-Glc^iii^(1→4)β-d-Glc^ii^(1→3)-d-Glc^i^], with partial assignment through ^13^C NMR (101 MHz, D_2_O) δ 102.94 (C-1^iii^ β), 102.69 (C-1^ii^ β), 102.60 (C-1^ii^ α), 95.68 (C-1^i^ β), 92.03 (C-1^i^ α), 84.52 (C-3^i^ β), 82.26 (C-3^i^ α), 78.29 (C-4^ii^ β), 75.57, 75.36, 74.86, 74.83, 74.21, 73.82, 73.20, 73.17, 72.51, 71.22, 71.04, 70.96, 68.57, 68.12, 68.08, 61.05, 60.73, 60.57, 60.06.

**Fig. 3.**
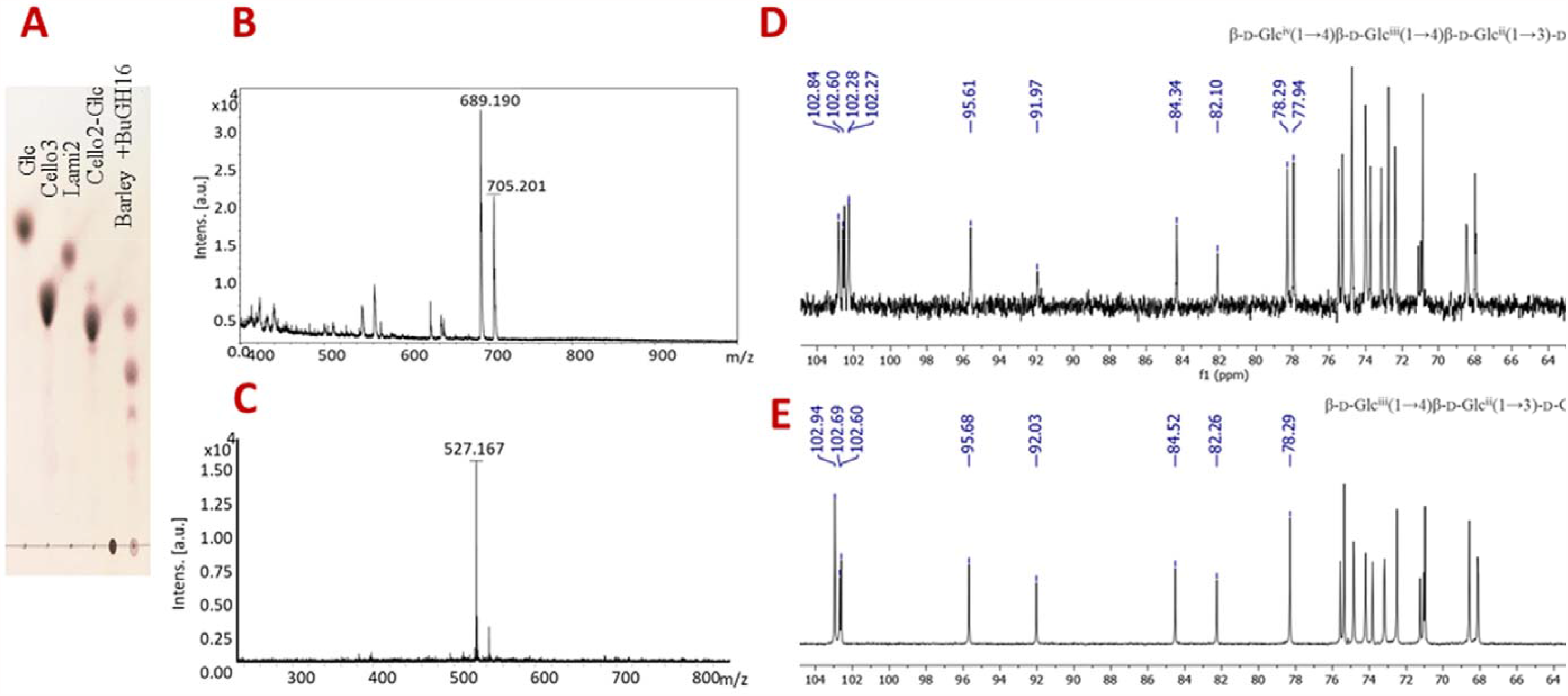
Hydrolysis of barley-β-glucan and characterization of its products. (**A**) The *Bu*GH16 of *Bacteroides uniformis* JCM 13288^T^ was exploited for producing mixed linkage oligosaccharides from barley β glucan. After enzymatic reaction, products were purified from Toyopearl S40 resin containing column, and (**B and C**) MALDI-TOF mass spectrum analysis was performed on both purified products. All masses were observed with sodium attached [M+ Na]^+^. (**D and E**) Purified mixed linkage oligosaccharides were characterized by ^13^C-DEPT-NMR.

### Determination of substrate specificity of BuGH3_MLG_

The BuGH3_MLG_ was initially screened to pinpoint specific activity against a variety of chromogenic pNP glycosides. Assays highlighted that *Bu*GH3_MLG_ can cleave β-1-3 and β-1-4 linked glucosides including a pNP-GlcNAc. Cleaving of pNP-Glc confirmed that it acts as exo-β glycosidase. *Bu*GH3_MLG_ exhibited slightly higher Kcat/*K*m for pNP-Glc as compared to pNP-cellobioside and pNP-laminaribioside (Table 2 and Fig. S1). The pNP-GlcNAc was a poor substrate with a 5.25, 4.6, 3.6 fold lower Kcat/*K*m than pNP-Glc, pNP-cellobioside and pNP-laminaribioside respectively. It did not hydrolyze pNP-rhamnopyranoside, pNP-galactopyranoside, and pNP-mannopyranoside (Table 2 and Fig. S1). Subsequently, enzymatic kinetics were carried out using PAHBAH reducing sugar assay on different gluco-oligosaccharides having β-1-3 and β-1-4 linkages. Kcat/Km was decreasing as degree of polymerization of gluco-oligosaccharides increases (Fig. 4F). Trends of kinetics parameters are found to be congruent with *Bo*GH3_MLG_ (Tamura, *et al*., 2017).

**Fig. 4.**
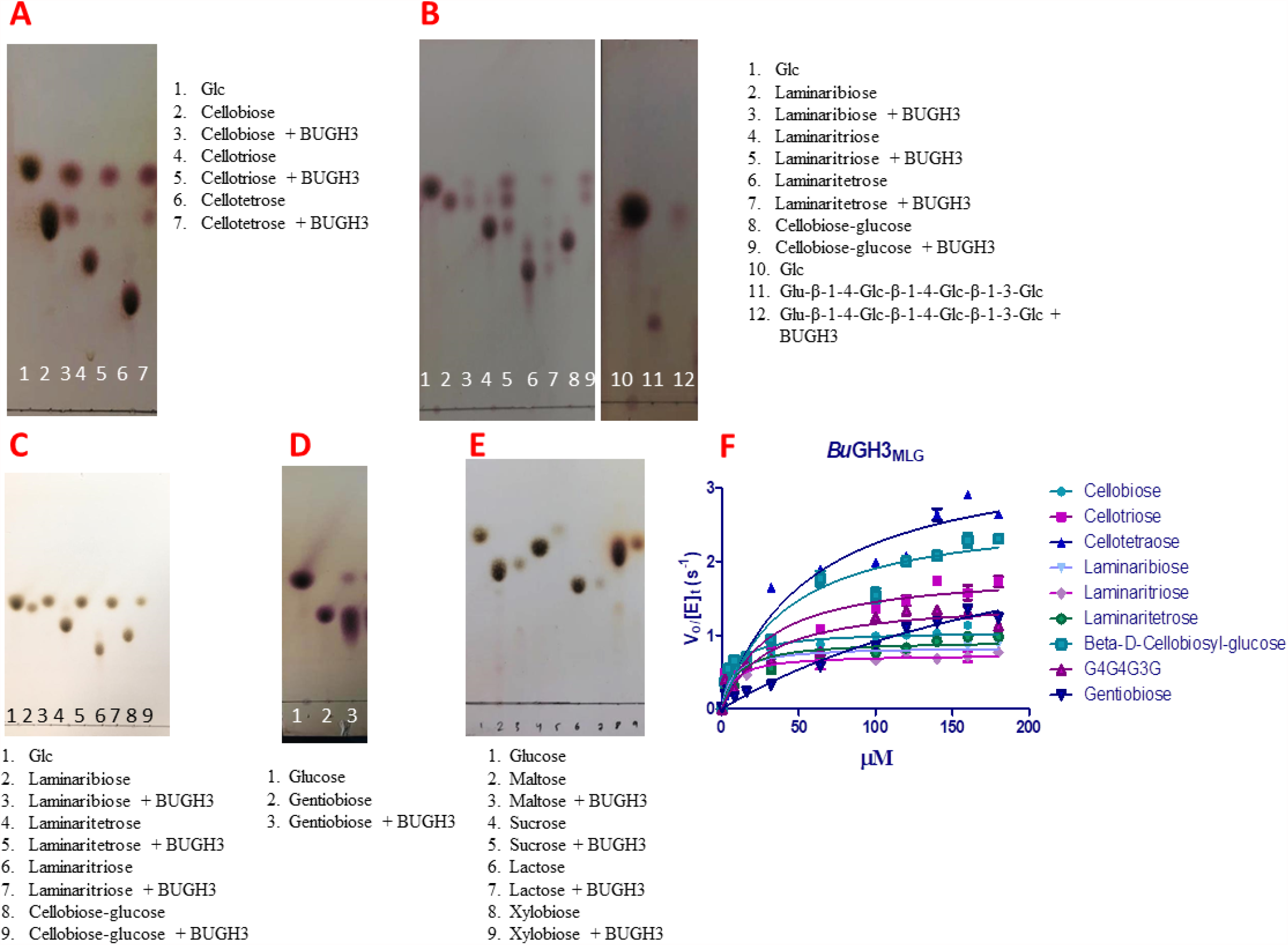
Hydrolysis of different oligosaccharides by the *Bu*GH3_MLG_ was evaluated by thin layer chromatography (TLC). TLC analysis reveals degradation pattern of cello-oligosaccharides (**A**), and laminari- and mixed linkage-oligosaccharides (**B**) after 4 h of enzymatic incubation. (**C**) TLC analysis reveals degradation pattern of laminari- and mixed linkage-oligosaccharides after overnight of enzymatic incubation. (**D**) TLC analysis reveals degradation pattern of gentiobiose, and (**E**) maltose, sucrose, lactose and xylobiose after 4 h of enzymatic incubation. TLC analysis was carried out on Silica Gel 60 F254 (Merck) and visualized by spraying TLC with 5% H_2_SO_4_ in ethanol followed by charring. (**F**) Michaelis-Menten plot demonstrating the enzyme kinetic with different substrates using PAHBAH reducing sugar assay. Kinetic curve was analyzed on GraphPad Prism.

Assessment of *Bu*GH3_MLG_ with gluco-oligosaccharides revealed that it hydrolyzed the used degree of polymerization of β-1-3 and β-1-4 linked glucosides within 4 h of incubation (Fig. 3A and B). The *Bu*GH3_MLG_ hydrolyzed β-1-3 and β-1-4 linked glucosides in an exo-acting manner from non-reducing end where the trisaccharide was first converted into disaccharide and then into monosaccharide (Fig. 3A, B and C). Interestingly, it also hydrolyzed gentiobiose (β-1-6) and very poorly sucrose. Normally gentiobiose (β-1-6 linked disaccharide) is generated during enzymatic hydrolysis of laminarin (Becker, *et al*., 2017) and *Bu*GH3_MLG_ was highly expressed when bacterium grew on minimal medium containing laminarin, suggesting that it plays a role in hydrolyzing gentiobiose (Fig. 1). In fact after 4 h of incubation, the enzymatic hydrolysis product was run on TLC, a spot correspondingly to standard glucose was noted. However, a LUL associated *Bu*GH3 of the *B. uniformis* JCM 5828 hydrolyses gentiobiose at slower rate and produces glucose when overnight incubation of reaction run on TLC (Singh, *et al*., 2020). The G4G3G and G4G4G3G were produced in the laboratory by the *Bu*GH16 and were then screened with BuGH3_MLG_. Substrate specificity demonstrates that G4G4G3G is first converted into G4G3G and then both compounds break down into glucose via laminaribiose as an intermediate (Fig. 4).

The *Bu*GH3_MLG_ showed 98% sequence homology to a gene (gene id: 2916276424) present in the genome of *B. uniformis* JCM 5828 when analyzed through the integrated microbial genomes and metagenomes (IMG/M) server. It highlights that the gene may play some role in BβG assimilation in case of *B. uniformis* JCM 5828. Specific activities of the *Bu*GH3_MLG_ are congruent with an enzyme *Bo*GH3_MLG_ of *B. ovatus* ATCC8483. The *Bo*GH3_MLG_ is a part of mixed – linkage BβG locus that acts on β-1-3, β-1-4 and β-1-6 (Tamura et al. 2017). Given the presence of β-1-3 and β-1-4 linkages in BβG and both enzymes showing comparable activities for hydrolyzing of a variety of gluco-oligosaccharide substrates, it can be concluded that the *Bu*GH3_MLG_ plays a very important role in degrading BβG.

Broader substrate specificity was reported from *Thermotoga neapolitana* and *Hypocrea jecorina* that showed activity on β-1-2, β-1-3, β-1-4 and β-1-6 linked oligosaccharide (Pozzo, *et al*., 2010, Karkehabadi, *et al*., 2014). The GH3 of barley exhibits narrower specificity to β-1-3 and β-1-4 linked glucan (Varghese, *et al*., 1999). Similar to barley GH3, xyloglucan utilizing PUL associated BoGH3A and BoGH3B from the *B. ovatus* displayed narrower specificity to β-1-4 linked gluco-oligosaccharides (Larsbrink, *et al*., 2014). Crystal structural information of these GH3 suggests that narrow and broader catalytic functionality associates with narrowing and more open active site architecture of the enzyme respectively (Hemsworth, *et al*., 2016). Much understanding about accommodation of diverse linkages gluco-oligosaccharides including mixed glucans can be obtained from the crystal structure of *Bu*GH3_MLG_; however, it is beyond the scope of the current study.

### Primary structural information and phylogeny analysis

GH3 is a large and divergent enzyme family with considerable differences in known substrate specificity, enzymatic hydrolyzing potential and active site architecture of amino acid residues. Phyre2 was used to obtain sequence identify percentage with resolved crystal structural proteins. Only 37% sequence identity with 100 % confidence matched to *Streptomyces venezuelae* (PDB number: 4I3G) and *Kluyveromyces marxianus* (PDB code 3AC0). The specific activity of GH3 of *K. marxianus* is restricted to disaccharides and it drastically decreases when trisaccharides were used (Yoshida, *et al*., 2010). GH3 of the *S. venezuelae* was reported to be involved in activating macrolide and was only tested with pNP-Glc (Zmudka, *et al*., 2013). *Bu*GH3_MLG_ showed substrate specificity similarly to *T. neapolitana*; however, they matched with only 32% sequence homology.

Most relevant resolved crystal structural protein sequences were downloaded from the RCSB-PDB database and other relevant enzymatic sequences were obtained from the NCBI for developing an evolutionary phylogenetic tree. In the phylogenetic tree, *Bu*GH3_MLG_ forms a cluster with a group of enzymes that display diverse substrate specificities (Fig. 5). It is shown more closely with BoGH3_MLG,_ where both share 72% sequence similarity. *Bo*GH3_MLG_ is also known to be active on β-1-3, β-1-4 and β-1-6 linked oligo-glucans (Tamura, *et al*., 2017). The tree suggests that *Bu*GH3_MLG_ and *Bo*GH3_MLG_ both evolutionary concomitantly developed to act on mixed linkage glucans. It can be predicted that in the course of evolution, GH3_MLG_ was maintained within the cluster whereas the *Bu*GH3_MLG_ could have been taken by horizontal gene transfer mechanism from other bacteria. *Bu*GH3_MLG_ is formed a distinct clade with *Bu*GH3_LM_, suggesting that both enzymes are evolved to act on different linked sugars with variable substrate specificities. *Bu*GH3_LM_ is a part of LUL of both *B. uniformis* strains and is highly active against β-1-3 linked oligosaccharides. Based on the above analyses, structure-based primary sequence alignment was performed on *Bu*GH3_MLG,_ *Bo*GH3_MLG_, and BT3314 (GH3) with structurally characterized GH3 β-glycosidase (PDB number: 2×41, *Thermotoga neapolitana*). The analysis again suggested more similarity between *Bu*GH3_MLG,_ *Bo*GH3_MLG_, and BT3314. Given the low sequence similarity with known resolved crystal structural protein (2×41), we could not be able to obtain secondary structural information of *Bu*GH3_MLG_. Nevertheless, structure-based sequence alignment predicted that in the absence of Trp-453 at the entrance site for ligand (Fig. S2), enzymes tend to show broad range of catalytic activity on β-1-2, β-1-3, β-1-4 and β-1-6 linked oligosaccharides. Notwithstanding, it warrants further investigation on the protein through the crystal structure.

**Fig. 5.**
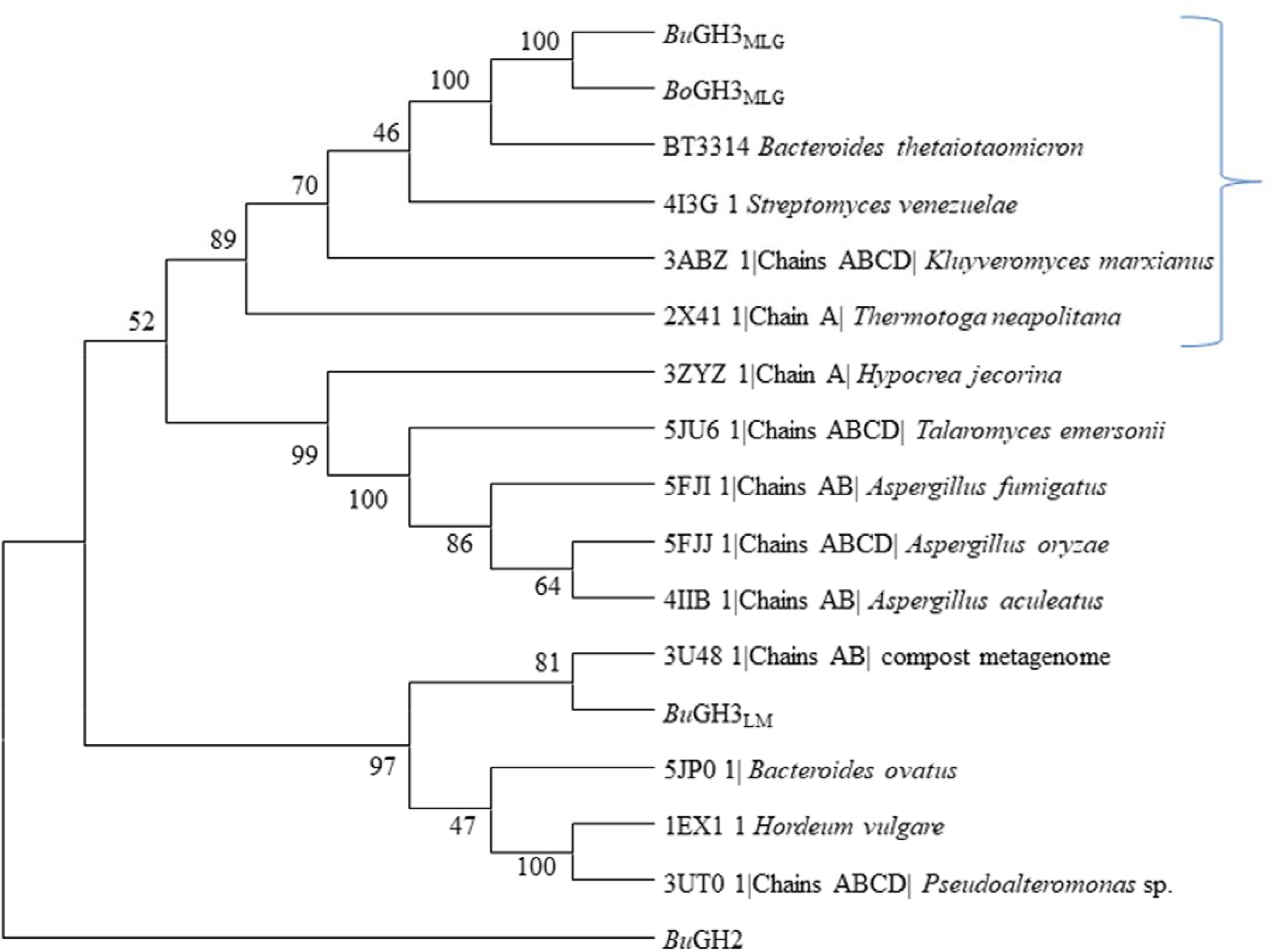
Neighbor-Joining phylogeny tree (Saitou & Nei, 1987) of relevant experimentally confirmed and resolved crystal structural GH3-glucosidases. The ClustalW algorithm was applied for amino acid sequence alignment and the tree was constructed with Poisson correction method where the bootstrap test (500 replicates) values are shown next to the branches (Felsenstein, 1985). Evolutionary analyses were conducted in MEGA X (Kumar, *et al*., 2018). *Bu*GH2 was used as an outgroup in the tree.

In both *B. uniformis* strains, LUL associated enzyme (BuGH16) breaks down barley, laminarin and lentinan, and produces mixed linkage (G4G4G3G and G4G3G), β-1-3 and β-1-6 linked oligosaccharides (Fig. 6 A). It was previously observed that pustulan-PUL in *B. uniformis* lacks exo-acting enzyme (GH3) in periplasmic space (Fig. 6 B). The pustulan-PUL associated *Bu*GH30 generates β-1-6 linked oligosaccharides, which are imported into the periplasm through the SusCD complex (Fig. 6 C). *Bu*GH3_LM_ associated with LUL was shown to be highly active on β*-*1, 3-linked oligosaccharides as compared to β*-*1, 4 and β*-*1, 6 -linked oligosaccharides, and hydrolyzed them into glucose before the oligosaccharides reach the cytoplasm via the Major Facilitator Superfamily (MFS) transporter (Dejean, *et al*., 2020, Singh, *et al*., 2020). A paucity of bespoke enzyme in PUL of laminarin and pustulan utilization can efficiently hydrolyse mixed-linkage and β-1-6 linked oligosaccharides generated from BβG and pustulan in the periplasm respectively (Dejean, *et al*., 2020, Singh, *et al*., 2020), the *Bu*GH3_MLG_ is most likely to take central role in degrading these oligosaccharides in the periplasm.

**Fig. 6.**
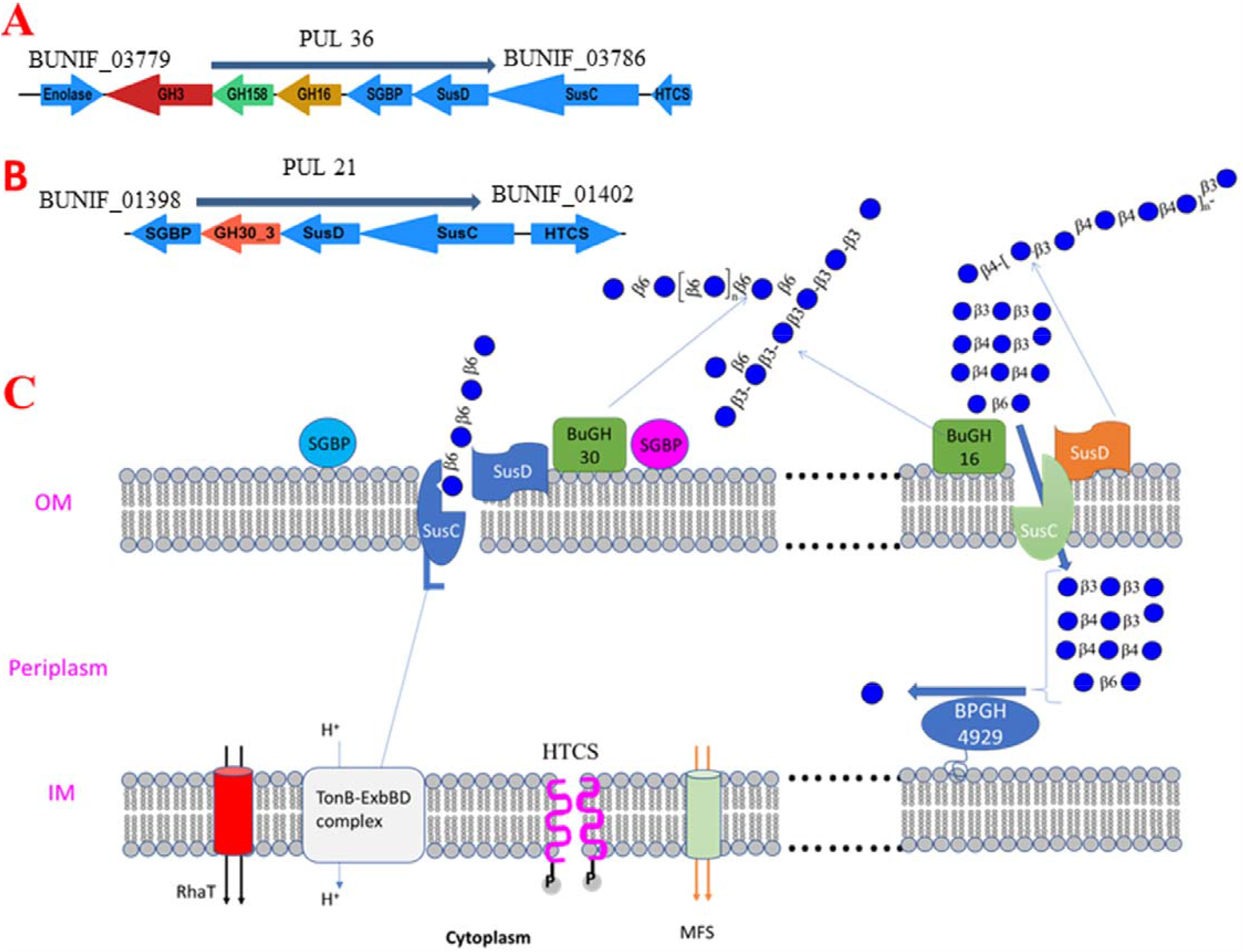
Possible role of the *Bu*GH3_MLG_ for utilizing oligosaccharides generated from enzymatic degradation of laminarin, pustulan and barley *β* glucan (BβG). Two PULs that utilize laminarin (PUL 36) and pustulan (PUL 21) were recently characterized (Singh, *et al*., 2020). In both strains, laminarin PUL (LUL) associated *Bu*GH16 breaks down BβG, laminarin and lentinan, and produces mixed linkages (G4G4G3G and G4G3G), β-1-3 and β-1-6 linked oligosaccharides respectively. Whereas pustulan PUL associated *Bu*GH30 breaks down longer chain β-1-6 linked glucan and generates β-1-6 linked oligosaccharides (Fig. 6 A). It was previously observed that pustulan PUL in *B. uniformis* lacks exo-acting enzyme (GH3) in periplasm (Fig. 6 B). These generated oligosaccharides import into the periplasm through the SusCD complex (Fig. 6 C). However, both PULs are paucity of bespoke enzymes that can hydrolyse mixed-linkage and β-1-6 linked oligosaccharides generated from BβG and pustulan respectively (Dejean, *et al*., 2020, Singh, *et al*., 2020). Thus, *Bu*GH3_MLG_ is most likely to take place central roles in degrading these oligosaccharides in the periplasm and allows bacteria to grow on these β glucans.

## Conclusion

Human genome has a bereft of genes encoding CAZymes and is not able to hydrolyze dietary fiber presence in our diet, which rationally emphasises on the importance of HGM (Tasse, *et al*., 2010, El Kaoutari, *et al*., 2013). The HGM has shown a tremendous capacity to dissect major dietary fiber (including xyloglucan, mannan and pectin) into their monomeric forms (Singh, 2019), and their metabolic products greatly impact well-being of the tissues in direct and indirect fashions (Lattimer & Haub, 2010). Furthermore, system biology has untangled many more functions of human gut bacteria to control metabolic diseases including irritable bowel diseases and diabetes (Koropatkin, *et al*., 2012). Omics approaches undoubtedly generate a huge number of data where diverse functional techniques have shown their importance to connect all of those available data and bring down to a reasonable form so as to understand adaption of each strain of human gut bacteria (Singh, *et al*., 2017). β*-*glucans are major sources of human diets and previous findings demonstrated that how *Bacteroides* utilize them through diverse molecular mechanisms (Larsbrink, *et al*., 2014, Ndeh, *et al*., 2017, Luis, *et al*., 2018, Foley, *et al*., 2019). A complete set of genes in the form of PUL has been observed to digest a specific type of β*-*glucan in different *Bacteroides*. It was also observed that LULs in the *B. uniformis* are able to hydrolyze mixed-linkage polysaccharides; even though it is lacking the bespoke exo-acting enzyme in the periplasm (Singh, *et al*., 2020) and is yet to be identify.

The essence of this study is that we identified a missing link, *Bu*GH3_MLG_ that acts on oligosaccharide generated by *Bu*GH16 and BuGH30. *Bu*GH3_MLG_ is found to hydrolyze β-1-3, β-1-4 and β-1-6 linked oligosaccharides including mixed linkage glucans (G4G4G3G and G4G3G). It is however not clear why *B. uniformis* adapted a unique strategy for utilizing mixed-linkage glucans in which the outer membrane-associated endo-acting enzyme is a part of PUL and an exo-acting enzyme in periplasm is not present within the PUL. Presumably, this strategy might have been adapted to avoid direct competition among *Bacteroides* by feeding on mixed linkage glucans.

## Acknowledgements

R. P. Singh would like to thank the Department of Biotechnology, India for providing the Ramalingaswami Re-entry Fellowship. The Executive Director of NABI is greatly appreciated for his continuous support for providing infrastructural facilities.

## Author contributions

RPS: Supervision, Conceptualization, Methodology, Data curation, Software, Investigation, Validation, Writing-Reviewing and Editing. RT: Investigation and Methodology. GK: Investigation and Methodology.

---

**Table 1:** Kinetics parameters for cleavage of various products by BuGH3_MLG_

| Substrate | $K_m$ (mM) | $K_{cat}$ (s<sup>-1</sup>) | $K_{cat}/K_m$ (s<sup>-1</sup> mM<sup>-1</sup>) |
| --- | --- | --- | --- |
| p-Nitrophenyl-β-D-glucopyranoside | 0.53 ± 0.12 | 17.8 ± 0.74 | 33.58 |
| p-Nitrophenyl β-D-laminaribioside | 0.88 ± 0.16 | 20.23 ± 0.63 | 23 |
| p-Nitrophenyl β-D-cellobioside | 0.64 ± 0.09 | 18.88 ± 0.47 | 29.5 |
| p-nitrophenyl-N-acetyl glucosaminide | 2.9 ± 0.7 | 18.5 | 6.4 |
| p-Nitrophenyl-β-D-galactopyranoside | ND | ND | ND |
| p-Nitrophenyl β-D-rhamnopyranoside | ND | ND | ND |
| 4-Nitrophenyl β- D – mannopyranoside | ND | ND | ND |
| Substrate | $K_m$ (μM) | $K_{cat}$ (s-1) | $K_{cat}/K_m$ (s<sup>-1</sup> μM<sup>-1</sup>)<br>PAHBAH assay |
| Cellobiose | 7.4 ± 1.24 | 10.56 ± 0.24 | 1.42 |
| Cellotriose | 26.14 ± 5.5 | 18 ± 0.5 | 0.69 |
| Cellotetraose | 56.9 ± 9 | 35.1 ± 0.8 | 0.29 |
| Laminaribiose | 7.1 ± 1.2 | 8.5 ± 0.2 | 1.2 |
| Laminaritriose | 7.5 ± 0.5 | 7.4 ± 0.17 | 1 |
| Laminaritetrose | 9.9 ± 1.1 | 9.3 ± 0.26 | 1 |
| β-D-cellobiosyl-glucose | 41.31 ± 7 | 26.9 ± 0.64 | 0.66 |
| G4G4G3G | 28.2 ± 6.2 | 14.83 ± 0.4 | 0.53 |
| Gentiobiose | 241.2 ± 16.2 | 39.5 ± 1 | 0.16 |
| Sucrose | ND | ND | ND |

The value of blank was subtracted from value of test value in PAHBAH reducing sugar assay. G4G4G3G: Glu-β-1-4-Glc-β-1-4-Glc-β-1-3-Glc. Given the low activity on sucrose, we could not determine kinetic parameters.

