## Supplementary File 1 for "Human gut Bacteroides uniformis utilizes mixed linked β- glucans via an alternative strategy"

**Table S1: Primers are used for RT-qPCR and cloning**

| **Name** | **Type** | **Sequence** |
| --- | --- | --- |
| RT_00608 | Forward primer | CGGTGGTAGTTCTTCCCTGA |
|  | Reverse primer | AGATTCTGTCCGGTCACCAC |
| RT_04929 | Forward primer | TCAAGCACTATGCGCTGAAC |
|  | Reverse primer | GGCCGCGAATCTTATTGTAA |
| RT_04508 | Forward primer | TTCCAATCTGAAACCGGAAC |
|  | Reverse primer | GATTCGCCCAAGAGAATGTC |
| Cloning of 04929 | Forward primer | CAGCAAATGGGTCGCGGATCCCAGACACCGGTCTATATGGACGA |
|  | Reverse primer | GTGGTGGTGGTGGTGCTCGAGTTATTGCATTTCGAACGAAACTACAC |

**Table S2:** The effect of metal ion on the activity of BUGH3

| **Additive** | **Concentration (mM)** | **% Relative activity** |
| --- | --- | --- |
| HEPES | 0 | 100 |
| MnCl_2_ | 2 | 78 ±2 |
| MgCl_2_ | 2 | 47 ± 5 |
| CaCl_2_ | 2 | 81 ±14 |
| FeSO_4_ | 2 | 86 ±10 |
| MgSO_4_ | 2 | 90 ±11 |
| LiBr | 2 | 100 ±16 |
| NaNO_3_ | 2 | 90 ±13 |
| NH_4_Cl | 2 | 87 ±11 |
| KCl | 2 | 95 ±16 |
| CoCl_2_ | 2 | 86 ±12 |
| NiSO_4_ | 2 | 86 ±12 |
| NaCl | 2 | 87 ±12 |
| NaCl | 5 | 131 ±13 |
| NaCl | 10 | 151 ±16 |
| Urea | 2 | 53 ±2 |
| DTT | 2 | 54 ±2 |
| EDTA | 2 | 64 ±13 |

± indicates the deviation of value from mean (OD) of the three replicates

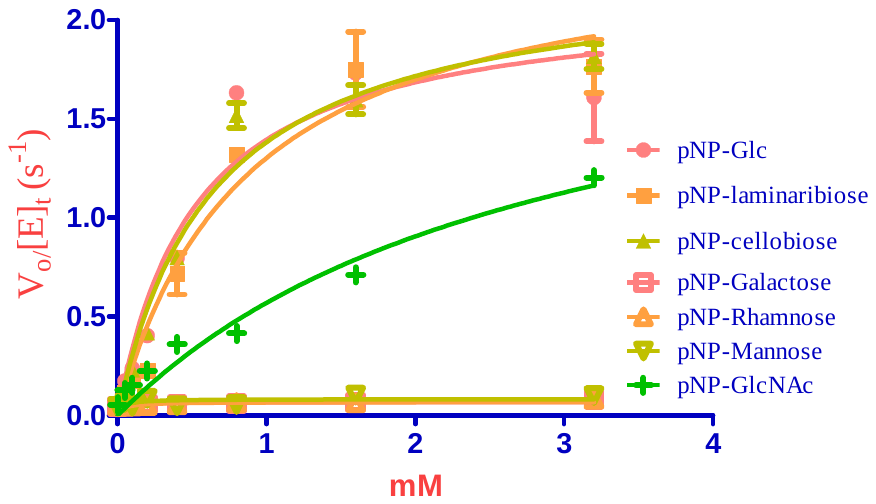

**Fig S1.** **Michaelis-Menten plot demonstrating the enzyme kinetic with different substrates**. p- Nitrophenyl-β-D-glucopyranoside (pNP-Glc), p-nitrophenyl β-D-laminaribioside (pNP-laminaribiose), p-nitrophenyl β-D-cellobioside (pNP-cellobiose), p-nitrophenyl-N-acetyl-glucosaminide (pNP-GlcNAc), p-nitrophenyl-β-D-galactopyranoside (pNP-galactose), p-nitrophenyl- β-D-rhamnopyranoside (pNP-rhamnose) and 4-nitrophenyl β- D –mannopyranoside (pNP-mannose). Kinetic curve was analyzed on GraphPad Prism.

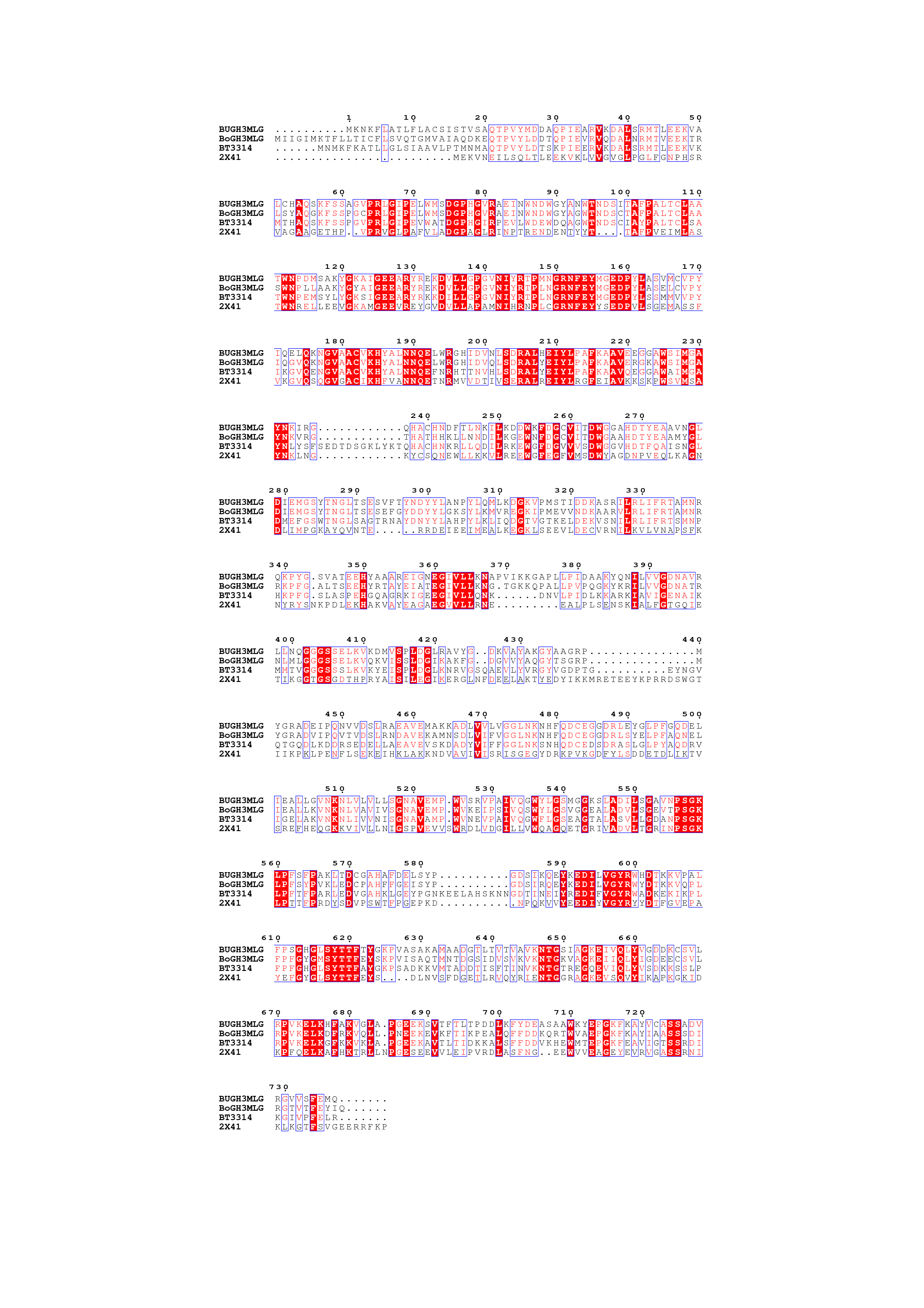

**Fig S2:** Structure-based sequence alignment of BuGH3_MLG,_ BoGH3_MLG_, and BT3314 (GH3) with structurally characterized GH3 β-glucosidases (PDB number: 2X41, *Thermotoga neapolitana*). Red highlights indicate unchanged positions and blue outlines indicate similar positions with variable amino acids. Alignment illustration performed with ESPript. 3.0 (Robert & Gouet, 2014).
